# Dopaminergic projections to the prefrontal cortex are critical for rapid threat avoidance learning

**DOI:** 10.1101/2024.05.02.592069

**Authors:** Zachary Zeidler, Marta Fernandez Gomez, Tanya A. Gupta, Meelan Shari, Scott A. Wilke, Laura A. DeNardo

## Abstract

To survive, animals must rapidly learn to avoid predictable threats. Such learning depends on detecting reliable cue-outcome relationships that efficiently drive behavioral adaptations. The medial prefrontal cortex (mPFC) integrates learned information about the environment to guide adaptive behaviors^1–7^ and is critical for threat avoidance^8–15^. However, as most studies have focused on well-learned threat avoidance strategies, the specific inputs that signal avoidability and drive rapid avoidance learning remain poorly understood. Dopamine (DA) inputs from the ventral tegmental area (VTA) potently modulate prefrontal function and are preferentially engaged by aversive stimuli^16–21^. Previous studies implicated VTA and prefrontal DA in aversive learning^22–25^, but their findings have been constrained by limited spatiotemporal resolution. Here, we used high-resolution tools to dissect the role of the VTA-mPFC DA circuit in rapid avoidance learning. Optogenetic suppression of VTA DA terminals in mPFC selectively impaired learning of a cued avoidance response, without affecting cue-shock association learning, reactive escape behaviors, or expression of previously learned avoidance. Using a fluorescent DA sensor, we observe rapid, event-locked DA activity that emerged transiently during the initiation of learning. Increased DA encoded aversive outcomes and their predictive cues, while decreased DA encoded their omission and predicted how quickly mice learned to avoid. In yoked mice lacking control over shock omission, these dynamics were largely absent. Together, these findings demonstrate that the VTA-mPFC DA circuit is necessary for rapid acquisition of proactive avoidance behaviors and reveal transient event-related DA signals that underlie this form of learning.

## RESULTS AND DISCUSSION

### VTA-mPFC DA terminal activity is required for avoidance learning but not for associating a cue with an aversive outcome

While mPFC plays a key role in threat avoidance, the inputs that signal avoidability and mediate learning remain poorly understood. Previous studies implicated VTA-mPFC DA circuit activity in aspects of fear and safety learning^19,22,25–30^, but the role of these projections in learning to avoid threats remains unknown. To investigate this, we trained mice in platform-mediated avoidance (PMA)^10,13^, where they first learn that a tone predicts a foot shock, and then learn to avoid the shock by navigating to a safety platform. A reward port on the opposite side of the chamber forces mice to choose between safety and obtaining reward (Fig. 1A-C). To test the contribution of DAergic projections to mPFC in learning, we expressed the inhibitory opsin Jaws in the VTA of TH-Cre mice and inhibited axon terminals in mPFC throughout the tone and shock period (Fig. 1D-G). We targeted the prelimbic (PL) subregion of mPFC, which has an established role in threat avoidance^42,43^. Viral expression and optic fiber targeting were confirmed with histology (Fig. S1A,B). Jaws-expressing mice had significantly fewer successful avoidance trials compared to GFP controls, particularly on the first day and early in the second day of training (Fig. 1H). On days 1 and 2, Jaws mice also had significantly more escape trials, when they leapt to the platform after the onset of the shock (Fig. 1I), suggesting they failed to learn preemptive avoidance, but still understood the platform was safe. VTA-mPFC DA terminal inhibition did not affect freezing behavior during the tone (Fig. 1J), indicating that it did not affect learning about the cue-shock association. VTA-mPFC DA terminal inhibition also did not induce real-time place preference or aversion (Fig. S1C—F), indicating that activity in this pathway is not inherently rewarding or aversive.

**Figure 1.**
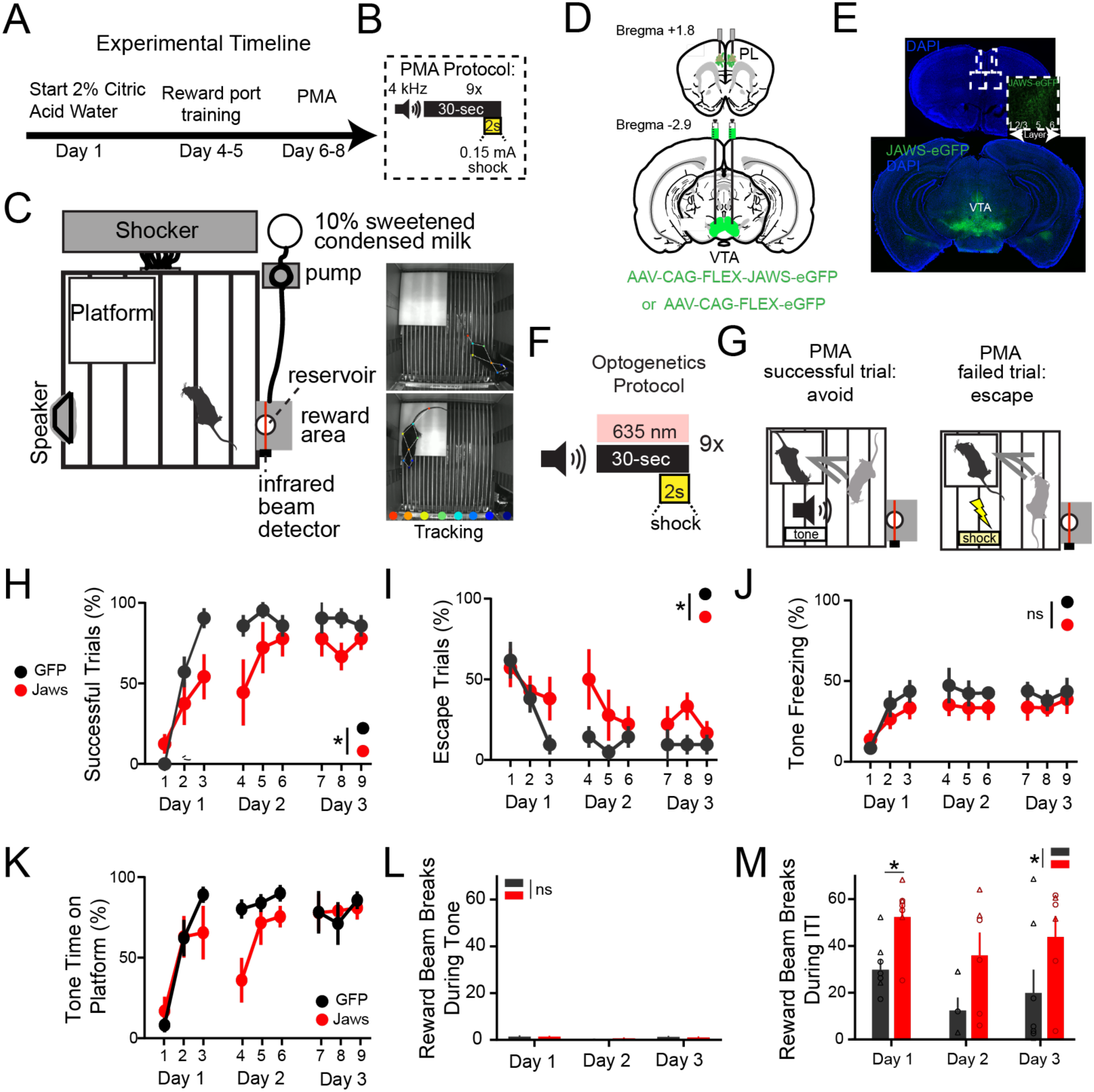
Optogenetic inhibition of VTA-mPFC DAergic axon terminals impairs avoidance learning but not tone-shock learning during PMA, See also Figure S1. A. Experimental timeline of PMA protocol. B. Behavioral training protocol. C. Schematic of PMA chamber (left) and frames from videos (right). D. Schematic of viral and fiberoptic targeting locations. E. Coronal sections from a representative brain showing Jaws-eGFP expression in VTA and bilateral fiber placement in PL. F. Optogenetic inhibition during PMA. 635nm laser light was presented coincidently with a 30 second 4kHz tone that co-terminated with a 2-second foot shock. G. Schematics showing a successful trial when mice preemptively avoided the shock vs. an escape trial when mice leaped to the platform after the shock began. H. Percent successful trials across days in GFP vs. Jaws mice (Mixed effects model F_time_(4.26, 49)=17, P<0.0001; F_opsin_(1, 12)=5.66, P=0.03; F_interaction_(8, 96) = 2.197, P=0.06; n=7 GFP, n=7-8 Jaws mice). I. Percent escape trials across days in GFP vs. Jaws mice (Mixed effects model F_time_(4,43, 49.86)=5.43, P=0.0007; F_opsin_(1, 12)=4.78, P=0.04; F_interaction_(8, 96)=1.04; P=0.4; n=7 GFP, n=7-8 Jaws mice). J. Percent time freezing during tone across days in GFP vs. Jaws mice (Mixed effects model F_time_(4.32, 46.98)=8.41, P<0.0001; F_opsin_(1, 12)=1.4, P=0.2; F_interaction_ (8, 96)=1.01; P=0.4; n=7 GFP, n=7-8 Jaws mice). K. Platform time during tone periods across days in GFP vs Jaws mice (Mixed effects model F_time_(4.2, 50) = 15, P<0.0001; F_opsin_(1, 12) = 0.9, P=0.3; F_interaction_(8, 96)=2.2, P=0.02; n=7 GFP, n=7-8 Jaws mice). L. Number of reward beam breaks during the tone period across days in GFP and Jaws mice (Mixed effects model F_time_(1.9, 32.1)=2.5, P=0.09; F_opsin_(1, 33)=5.66, P=0.9; F_interaction_(2, 33)=0.2, P=0.7; n=7 GFP, n=7-8 Jaws mice). M. Number of reward beam breaks during ITI across days in GFP and Jaws mice (Mixed effects model F_time_(1.9, 20.1) = 4.3, P=0.02; F_opsin_(1, 12)=8.2, P=0.01; F_interaction_(2, 21)=0.02, P=0.98; GFP vs Jaws Day 1 P=0.02; Day 2 P=0.2; Day 3 P=0.3, n=7 GFP, n=7-8 Jaws mice). *P<0.05, Graphs represent mean ± SEM. Circles denote males, triangles denote females.

Jaws mice tended to spend less time on the platform during tones (Fig. 1K), especially on the second day of training. Both groups showed minimal reward-seeking behavior during the tone, when avoidance behaviors are more adaptive (Fig. 1L). Yet during the inter-tone interval (ITI), opsin-expressing mice interacted with the reward port more than controls (Fig. 1M), suggesting that inhibiting VTA-mPFC DA terminals did not generally lead to reduced motivation to seek rewards. These findings indicate that VTA-mPFC DA projections are essential for cued avoidance learning, particularly during the early stages, but are not required for tone-shock learning or for reactive escape behaviors.

### mPFC DA signals evolve with PMA learning and predict future avoidance success

Previous experiments that used microdialysis reported elevations in mPFC DA during avoidance learning, but lacked the temporal precision required for a detailed analysis of how event-locked DA signals evolve during learning^24^. To directly observe mPFC DA dynamics during rapid avoidance learning, we recorded fluorescent signals using the DA sensor GRAB_DA2m_^31^ (GRABDA) during PMA (Fig. 2A,B; Fig. S2A). Consistent with the critical role of VTA-mPFC DA terminal activity early in learning (Fig. 1H,K), we observed the largest mPFC DA fluctuations on Day 1 of PMA (Fig. 2C). To understand how DA activity evolves across learning, we quantified mPFC DA levels and rates of change in our subsequent analysis (Fig. 2D) and examined relationships with PMA performance. Unsuccessful trials resulting in shocks initially evoked large DA responses that diminished with repeated exposures (Fig. 2E). On successful trials resulting in shock omissions, we observed decreases in mPFC DA activity that diminished with repeated avoids (Fig. 2F). Tone onset responses were smaller and did not consistently vary in magnitude as animals learned PMA (Fig. 2G).

**Figure 2.**
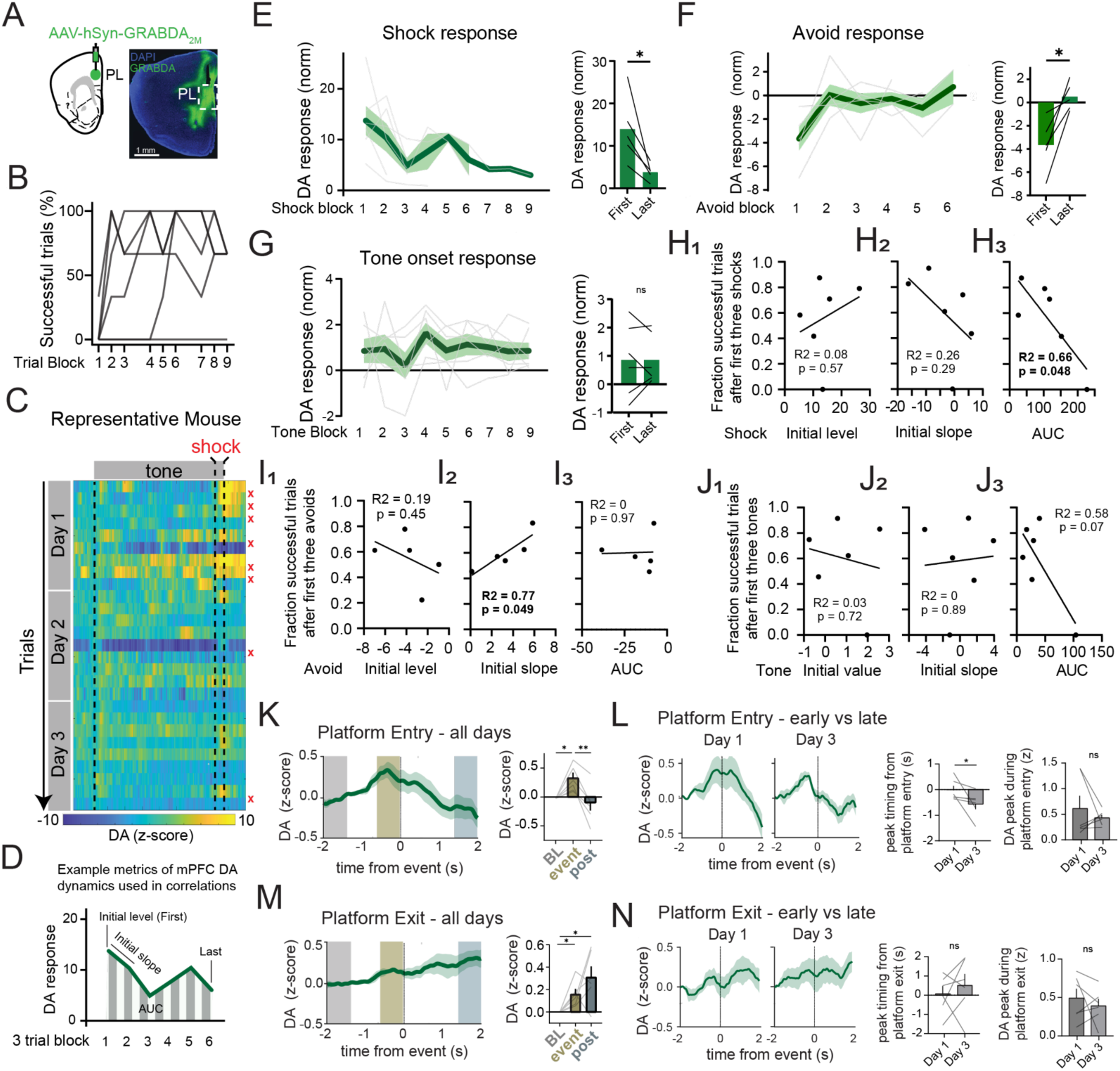
Prefrontal DA signals evolve with PMA learning and predict future avoidance success, See also Figures S2 and S3. A. Representative coronal section showing AAV-GRABDA_2m_ expression and fiber placement. B. Successful trials across time per animal (n=5 mice). C. z-scored DA fluorescence in a representative mouse. Red x marks shock trials. D. Illustration of metrics used to quantify mPFC DA dynamics and correlate with PMA performance. E. Left: mPFC DA signal during shocks across all sessions. Right: mean of first three shocks vs mean of last three shocks (paired t-test, P=0.03). F. Left: mPFC DA signal across successful avoids across all sessions. Right: mean DA during first three avoids vs mean DA during last three avoids (paired t-test, P=0.04). G. Left: DA signal tone onset across all sessions. Right: mean DA during first three tones vs mean DA during last three tones (paired t-test, P=0.9). H. Linear regression of successful trials after the first three shocks with mPFC DA activity during shocks (H_1_) initial DA level during block 1 (R^2^=0.08, P=0.57), (H_2_) slope between first two response blocks (R^2^=0.26, P=0.29), (H_3_) area under the curve (AUC) of all responses (R^2^=0.66, P=0.048). I. Linear regression of successful trials after the first three avoids with mPFC DA activity during successful avoids (I_1_) initial DA level during block 1 (R^2^=0.19, P=0,46), (I_2_) slope between first two avoid response blocks (R^2^=0.77, P=0.049), (I_3_) AUC of all responses (R^2^=0, P=0.97). J. Linear regression of successful trials with mPFC DA activity during tone onset (J_1_) initial DA level during block 1 (R^2^=0.03, P=0.72), (J_2_) slope between first two tone response blocks (R^2^=0, P=0.89), (J_3_) AUC of all responses (R^2^=0.58, P=0.07). K. Left: DA dynamics surrounding platform entry at baseline (BL), immediately preceding the behavior (event), or after the behavior (post) averaged across all platform entries on all days. Right: Quantification of signal changes (repeated measures one-way ANOVA F(1.4, 7.4)=10, P=0.01. BL vs. event P=0.02; event vs. post P=0.007; BL vs. post P= 0.7, Tukey’s multiple comparisons tests). L. Left: Comparison of early (Day 1) and late (Day 3) platform entry DA dynamics. Middle: Timing of signal peak from Day 1 and Day 3 (Wilcoxon signed rank test, P=0.031). Right: Peak of signal from Day 1 and Day 3 (Wilcoxon signed rank test, P=0.9). M. Same as K for platform exits (repeated measures one-way ANOVA F(1.2,6)=6, P=0.04. BL vs. event P=0.03. Event vs. post P=0.4. BL vs. post P=0.04, Tukey’s multiple comparisons tests). N. Same as L for platform exits. Middle: timing of peak signal from Day 1 to Day 3 (Wilcoxon signed rank test P=0.56). Right: peak of signal from Day 1 to Day 3 (Wilcoxon signed rank test P=0.68). *P<0.05, ** P<0.01. Graphs represent mean ± SEM.

We reasoned that both aversive shocks and their omissions may potentially drive PMA learning. We therefore investigated whether aspects of the mPFC DA activity during these events correlated with learning (Fig. 2D). DA can alter mPFC computations associated with learning^32^, so we also examined the relationships between tone-evoked DA activity and learning. We first focused on trial blocks one and two, where we observed the largest DA signals. Neither the initial level nor the initial rate of decrease (slope) in shock-evoked DA significantly correlated with PMA performance (Fig. 2H_1-2_). However, the area under the curve (AUC) for *all* shock blocks was significantly correlated with worse PMA performance (Fig. 2H_3_), suggesting that shocks continue to elicit large increases in DA when learning is still incomplete.

In contrast to shock trials, the AUC for DA activity on successful trials was not correlated with PMA performance (Fig 2I_3_). Neither was the initial magnitude of the negative deflection (Fig 2I_1_). However, the initial negative slope after shock omission was positively correlated with PMA performance (Fig 2I_2_), suggesting that a quickly diminishing response of mPFC DA may drive avoidance learning. Tone-evoked mPFC DA fluctuations were not significantly correlated with PMA performance (Fig. 2J_1-3_), suggesting that mPFC DA activity during trial outcomes may be more important for driving avoidance learning.

We also observed fluctuations in mPFC DA throughout tone presentations (Fig. 2C) which we hypothesized could be associated with behaviors important for avoidance learning. mPFC DA ramped up, peaking prior to platform entries and returning to baseline once mice reached safety (Fig. 2K). Interestingly, the magnitude of the mPFC DA peak did not change across learning, but the timing of the peak response shifted to precede platform entry on Day 3, once avoidance learning was complete (Fig. 2L). Conversely, mPFC DA increased following platform exits, remaining elevated as mice explored the shock grid (Fig. 2M). There was no change in the peak timing or magnitude across learning (Fig. 2N). We also examined DA during freezing behavior, which showed trend-level decreases prior to onset (Fig. S2D–F) and increases prior to offset of freezing (Fig. S2G–I). There were no significant changes in timing or magnitude of freezing-related DA activity across PMA learning (Fig. S2D,I). This suggests that avoidance learning selectively shapes mPFC DA activity related to danger-to-safety transitions, but not safety-to-danger transitions or reactive freezing.

Control experiments confirmed that GRABDA dynamics were not due to movement or motion artifacts. Movement velocity was uncorrelated with GRABDA signals (Fig. S3A,B) and fiber photometry recordings in GFP-expressing animals showed no shock or avoidance responses (Fig. S3C–F). Together, our findings indicate that mPFC DA encodes trial outcomes, conditioned cues, and threat-induced behaviors. Moreover, changes in both shock- and shock omission-, but not tone-evoked mPFA DA correlated with PMA performance, linking specific event-locked mPFC DA dynamics with avoidance learning. To further investigate these relationships, we next recorded mPFC DA in aversive learning paradigms that lacked avoidability, or in which shocks were sometimes omitted, but were not under behavioral control.

### mPFC DA activity encodes the predictability and avoidability of aversive events

Previous studies have reported increases in mPFC DA and VTA DA neuron activity during unexpected omission of an aversive stimulus^33,34^. However, we find that *decreases* in mPFC DA encode shock omissions when a behavior leads to successful avoidance (Fig. 2F, I_2_). We therefore designed control experiments to better understand how mPFC DA dynamics relate to the avoidability and predictability of aversive outcomes. First, we recorded mPFC DA activity in control mice with shocks and shock-omissions yoked to the individual mice experiencing PMA, but without a safety platform (Fig. 3A,B; Fig. S2B). Thus, both PMA and yoked groups experienced identical patterns of shocked and non-shocked trials, but in the yoked group there was no behavioral control over the outcome. Similar to PMA, mice exhibited increased freezing to the tone (Fig. 3C), and elevations in mPFC DA during shocks and tone onsets (Fig. 3D). However, unlike in PMA, shock-evoked DA in the yoked condition remained persistently elevated across trials (Fig. 3E). Strikingly, the dip in mPFC DA seen during early shock omissions in PMA was completely absent in yoked controls (Fig. 3F). This suggests that in PMA, negative DA signals encode behavioral control over avoidance rather than shock omission. In further contrast to PMA, DA levels during tone onset increased significantly over time (Fig. 3G). This suggests that tone-evoked mPFA DA increases when the relationships between cues, aversive outcomes, and behavioral strategies are uncertain.

**Figure 3.**
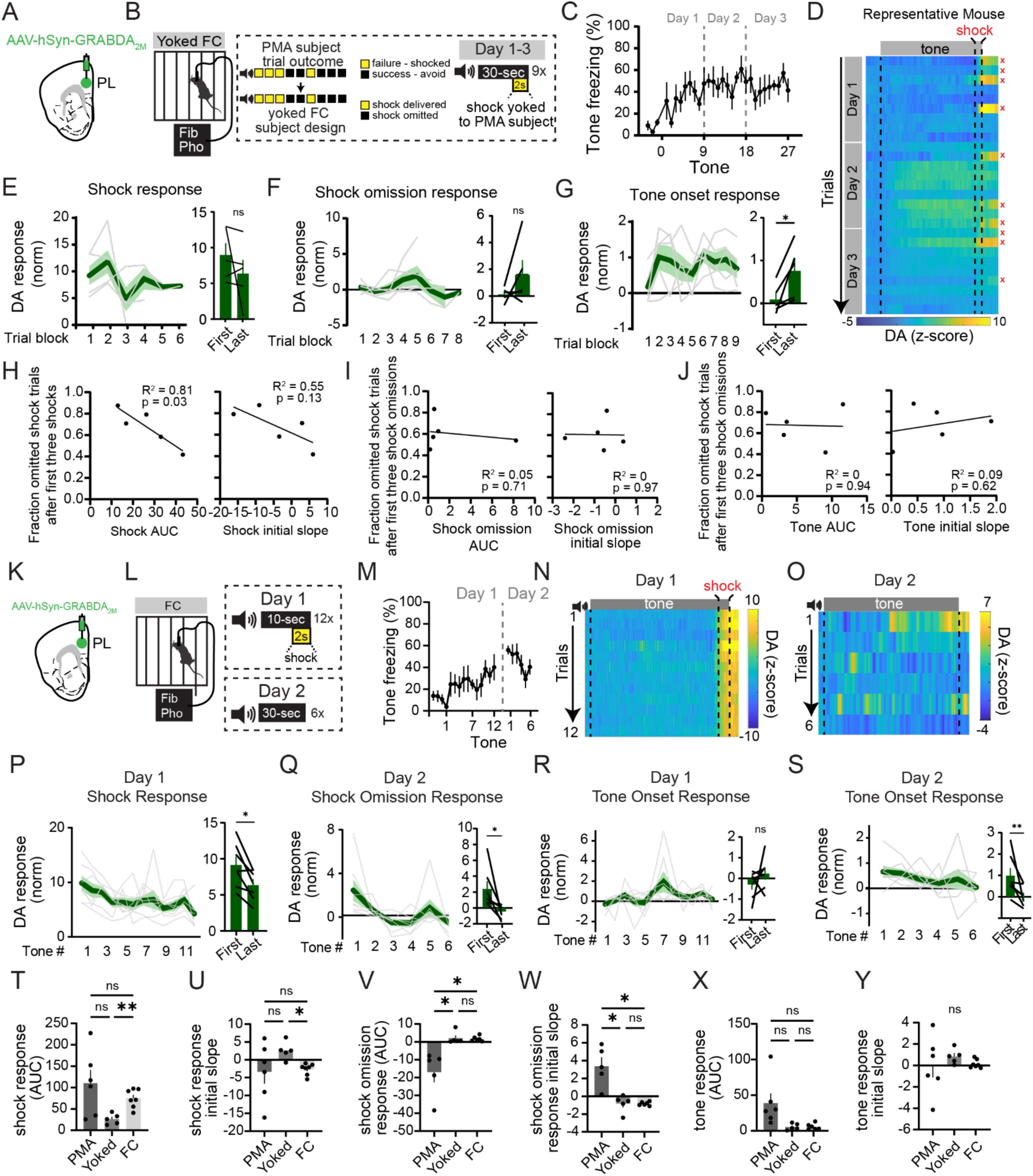
mPFC DA activity is sensitive to the avoidability and predictability of aversive outcomes, See also Figure S4. A. Schematic showing GRABDA recording in prelimbic (PL) cortex. B. Yoked experimental design: animals were placed in a fear conditioning (FC) arena and exposed to tones. Presence of a co-terminal shock was dependent on the matched trial outcome of a subject from PMA. C. Freezing during the tone across days (n=5 mice). D. Heatmap of GRABDA fluorescence (zscored) from representative mouse. E. Left: DA levels during shocks across time. Right: Comparison of DA during first vs. last shock block (paired t-test, P=0.3, n=5 mice). F. Left: DA levels during shock omission across time. Right: Comparison of DA during first vs. last shock omission block (paired t-test P=0.2, n=5 mice). G. Left: DA levels during tone onset across time. Right: Comparison of DA during first vs. last tone block (paired t-test P=0.03, n=5 mice). H. Linear regression of successful trials after the first shock block with (left) area under the curve (AUC) of shock responses (R^2^=0.81, P=0.03, n=5 mice) and (right) slope between first two shock response blocks (R^2^=0.55, P=0.13, n=5 mice). I. Linear regression of successful trials after the first avoid block with (left) area under the curve (AUC) of avoid responses (R^2^=0.05, P=0.71, n=5 mice) and (right) slope between first two avoid blocks (R^2^=0, P=0.97, n=5 mice). J. Linear regression of successful trials with (left) area under the curve (AUC) of tone responses (R^2^=0, P=0.94, n=5 mice) and (right) slope between first two tone response blocks (R^2^=0.09, P=0.62, n=5 mice). K. AAV and fiber placement for GRABDA fiber photometry in fear conditioning. L. Experimental design for fear conditioning recordings. M. Freezing during tone (n=6 mice). N. Heatmap from representative mouse showing GRABDA fluorescence during fear conditioning. O. Same as N for fear memory retrieval. P. Left: DA levels during shock during across the fear conditioning session. Right: Comparison of first to last shock response (paired t-test P=0.01, n=6 mice). Q. Left: DA levels during end of tone (shock omission) across the retrieval day session. Right: Comparison of first and last tone response (paired t-test P=0.03, n=6 mice). R. Left: DA levels during tone onset across the fear conditioning session. Right: Comparison of first and last tone response (paired t-test P=0.3, n=6 mice). S. Left: DA levels during tone onset across the fear memory retrieval session. Right: Comparison of first and last tone response (paired t-test P=0.008, n=6 mice). T. Comparing behavioral assays: shock response AUC (Welch’s ANOVA W(2,8.9)=13.4, P=0.002; PMA vs Yoked P=0.11; PMA vs FC P=0.63; Yoked vs FC P=0.001, Dunnet’s T3 multiple comparisons test). U. Comparing behavioral assays: shock response slope (Welch’s ANOVA W(2, 7.5)=7.8, P=0.014; PMA vs Yoked P=0.33; PMA vs FC P=0.99; Yoked vs FC P=0.017, Dunnet’s T3 multiple comparisons test). V. Comparing behavioral assays: avoid response AUC (Welch’s ANOVA W(2,4.6)=10.1, P=0.019. PMA vs Yoked P=0.025, PMA vs FC P=0.029, Yoked vs FC P=0.82, Dunnet’s T3 multiple comparisons test). W. Comparing behavioral assays: avoid response slope (Welch’s ANOVA W(2,5.5)=8, P=0.022. PMA vs Yoked P=0.029, PMA vs FC P=0.033, Yoked vs FC P=0.99). X. Comparing behavioral assays: tone onset response AUC (Welch’s ANOVA W(2,5.3)=5.5, P=0.048. PMA vs Yoked P=0.15, PMA vs FC P=0.15, Yoked vs FC P=0.99, Dunnet’s T3 multiple comparisons test). Y. Comparing behavioral assays: tone onset response slope (Welch’s ANOVA W(2,7.2)=1.8, P=0.2). * P<0.05, ** P<0.01, ns P>0.05. Graphs represent mean ± SEM.

Similar to our PMA recordings, we examined correlations between mPFC DA activity and shock-omission trials in the yoked group, focusing on the measures we observed to be correlated with PMA performance. The AUC for shock-evoked DA was negatively correlated with shock-omission trials (Fig. 3H) in a manner similar to PMA. This suggests that the total DA response across sessions reflects shock exposure rather than avoidance learning. Importantly, unlike in PMA, the initial slope of the DA response to early shock omissions and tones was not correlated with future shock outcomes (Fig. 3I,J). The findings indicate that the relationships observed during PMA for these same features are likely due to behavioral control over avoidability during learning, rather than as an indirect consequence of the pattern of shock omissions.

While our yoked task design maintained the tone-outcome sequence of PMA, it also made tone-shock relationships unpredictable. In PMA, this relationship is both predictable and avoidable. To examine the role of VTA-mPFC DA terminals and DA dynamics when mice learn about shocks that are predictable but unavoidable, we contrasted our previous results with cued fear conditioning. Inhibiting VTA-mPFC DA terminals during fear conditioning had no effect on freezing during conditioning or recall (Fig. S4). These results indicate that activity in the DAergic VTA-mPFC pathway is not required for learning tone-shock associations, even without an avoidance contingency.

We then recorded DA dynamics during one day of fear conditioning and during fear memory retrieval in which tones are presented without shocks (Fig. 3K-O; Fig. S3C). As in PMA, shocks evoked large increases in mPFC DA that steadily declined in amplitude with successive shocks (Fig. 3P). However, during retrieval there was initially a large *increase* in DA with the first shock omission that rapidly attenuated (Fig. 3Q). This contrasts with the *decrease* in DA seen with early shock omissions during PMA (Fig. 2F) and the lack of DA response to shock omission in yoked controls (Fig. 3F). Together these findings indicate that DA activity during shock omissions is distinctly modulated under conditions based on the avoidability and predictability of aversive outcomes. Tone onset responses during fear conditioning showed no consistent change (Fig. 3R), but during retrieval, they significantly decreased across repeated tone presentations (Fig. 3S), suggesting they may be associated with the behavioral significance of the tone.

To better understand how predictability and avoidability of aversive outcomes modulate mPFC DA, we directly compared responses to shock, shock-omission, and tone onset for all three groups. We only observed significant differences in the overall shock response (AUC) and rate of change in the shock response (slope) between the fear conditioned and yoked groups (Fig. 3T,U). This suggests that shock-evoked DA encodes predictability and may not play a role in avoidance learning. The overall shock omission response (AUC) on successful trials during PMA was larger and the initial slope was steeper compared to shock omissions for either yoked or fear conditioned groups (Fig. 3V,W). We observed no significant differences between the amplitude of mPFC DA activity at tone onset, or for the initial slope of the tone response (Fig. 3X,Y). Taken together, these findings suggest that, compared to mPFC DA activity during shocks, mPFC DA during shock omissions may play a more important role in learning to preemptively avoid threats. Moreover, because these dynamic activity patterns are transient, they may play a more important role in learning to avoid than in established avoidance.

### VTA-mPFC DA terminal activity is required for avoidance learning without motivational conflict, but not retrieval of a previously learned avoidance strategy

The mPFC is strongly modulated by motivational conflict^35–38^. Thus, we wondered whether the conflict presented between safety and reward-seeking was critical for the role of VTA-mPFC terminals in PMA. To investigate this, we trained mice in a version of PMA that lacked a competing reward (Fig. 4A,B). Similar to the mice trained in PMA with a motivational conflict (Fig. 1), this distinct group of opsin-expressing mice exhibited fewer successful avoidance trials and more shock-triggered escapes, but no change in tone-elicited freezing during PMA training (Fig. 4C). During a retrieval test the following day (without shocks or optogenetic inhibition), opsin-expressing mice continued to spend less time on the platform than controls despite similar freezing behavior (Fig. 4D). Thus, the cue-shock association remained intact, but they were unable to consolidate threat avoidance learning. Taken together, these data indicate that DAergic inputs to mPFC are required in a similar manner for avoidance learning, irrespective of the need to balance avoidance with reward-seeking behaviors.

**Figure 4.**
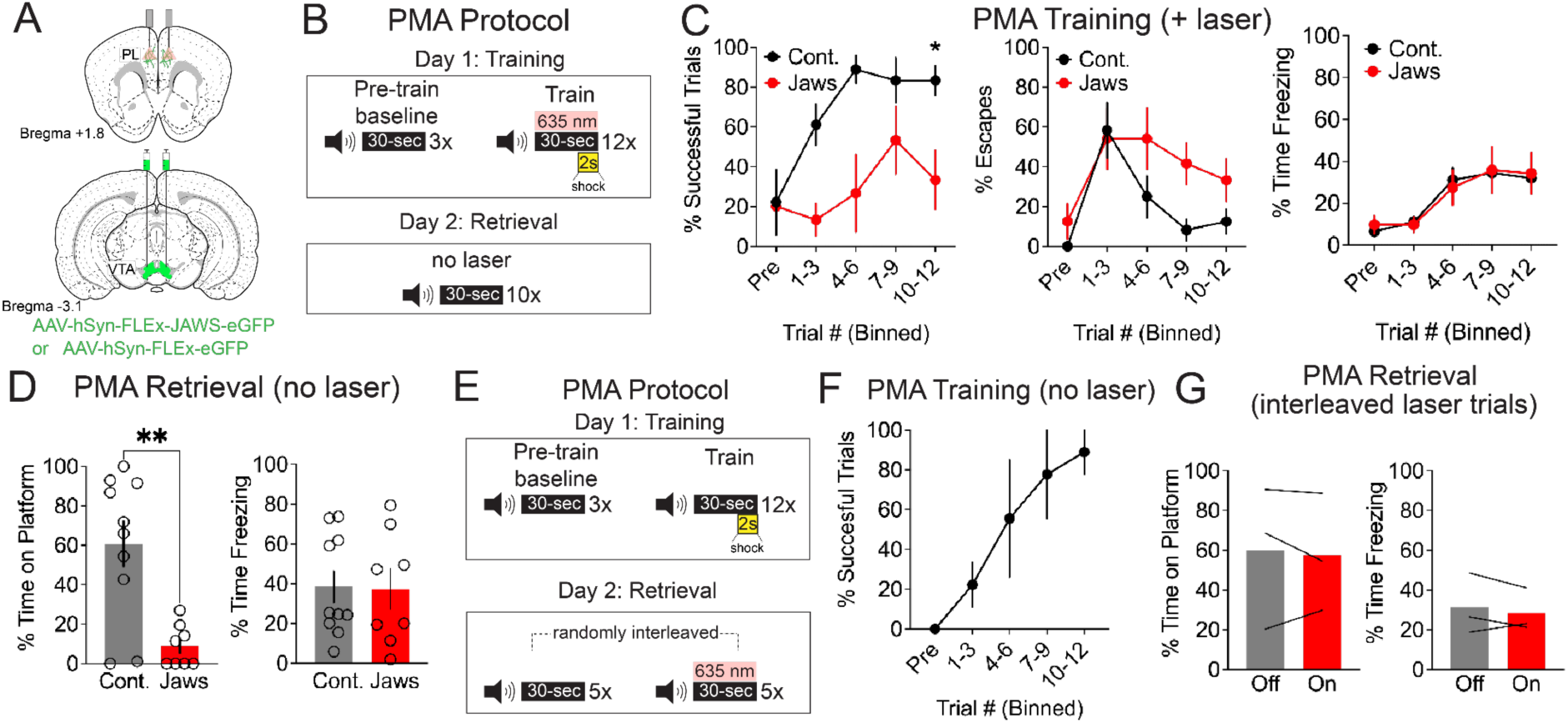
Optogenetic inhibition of VTA-mPFC DA terminals during PMA without motivational conflict and following learning. A. Schematic of viral and fiber optic targeting locations. B. Protocol for optogenetic inhibition during PMA training. On training day, 635nm laser light was presented coincidently with a 30 second 4kHz tone that co-terminated with a 2-second footshock. On retrieval day, mice were presented with 10 tones without shocks or laser. C. Left: Fraction of successful trials across days in control vs. Jaws mice (F_time_(4.323, 49.71)=16.66, P<0.0001; F_opsin_(1, 13)=5.092, P=0.04; F_interaction_(8, 92)=2.197, P=0.03; Mixed effects model, n=7 Jaws, n=8 control mice). Middle: Fraction of escape trials across days in control vs. Jaws mice (F_time_ (3.975, 44.71)=2.33, P=0.07; F_opsin_(1, 12)=1.68, P=0.22; F_interaction_ (8, 90)=3.09; P=0.004. Mixed effects model, n=7 Jaws, n=8 control mice). Right: Fraction time freezing during tone across days in control vs. Jaws mice (F_time_(4.39, 46.6)=8.39, P<0.0001; F_opsin_(1, 11)=0.60 P=0.46; F_interaction_(8, 85)=0.88; P=0.54. Mixed effects model, n=7 Jaws, n=7 control mice). D. Fraction time on platform and fraction of time freezing during tones on retrieval day (platform: P=0.002, freezing: P=0.6, two-tailed t-test, n=10 control, n=8 Jaws mice). E. Protocol for optogenetic inhibition during PMA retrieval. Mice are trained in PMA without laser inhibition. The next day, mice are presented with 10 randomly interleaved tones, half of which are paired with constant 635 nm light. F. Successful trials during PMA training (n=3 mice). G. Percent time on platform and percent time freezing during the tone on interleaved light-on and light-off trials (n=3 mice; Platform Wilcoxon signed-rank test P=0.75; Freezing Wilcoxon signed-rank test P=0.5). *P<0.05, **P<0.01. Graphs represent mean ± SEM.

Our data reveal a critical role for VTA-mPFC DA activity in avoidance learning, particularly when rapidly learning the implications of a successfully avoided aversive outcome (Fig. 1,2). However, we wondered whether VTA-mPFC DA terminal activity was selectively required for avoidance learning, or whether it is also required to retrieve a previously learned avoidance strategy. To address this question, we first trained a completely separate cohort of opsin-expressing mice in PMA (but without any optogenetic inhibition) (Fig. 4E,F). Then, during a retrieval session the following day, we interleaved trials with or without optogenetic inhibition during the 30s tone period (Fig. 4E). Inhibiting VTA-mPFC DA terminals had no effect for time on platform or freezing during the tone period (Fig. 4G). Thus, VTA-mPFC DA terminal activity is selectively required to link predictive cues with avoidance behaviors during rapid avoidance learning, but not for retrieving a previously learned avoidance strategy.

## Discussion

We demonstrate a critical role for VTA-mPFC DA activity in learning to avoid signaled threats. We reveal the event-locked patterns of mPFC DA activity and how they change as animals learn the relationships between cues and aversive outcomes, and the strategies required to avoid them. By comparing mPFC DA activity in three distinct, but related, aversive learning paradigms, we show how mPFC DA dynamically encodes predictive cues, aversive outcomes or their omission, and threat-related behaviors. Predictability influenced how quickly, and to what extent, DA signals diminished during subsequent presentations of aversive shocks. Avoidability was encoded in the direction and magnitude of DA activity during shock omissions. During PMA, when shocks were avoidable, we observed a negative deflection in mPFC DA coincident with successful avoidance during early learning. In contrast, when shock omissions were unpredictable (yoked controls), mPFC DA responses were minimal and exhibited no changes across a session. Moreover, during conditioned fear retrieval, we observed a positive deflection in mPFC DA during shock omissions that rapidly diminished. Optogenetic inhibition of VTA-mPFC DA terminal activity significantly slowed avoidance learning in PMA with or without a motivational conflict, but did not influence cue-shock learning or maintenance of learned avoidance. Our data underscore the necessity of DA for rapidly linking predictive cues to adaptive avoidance behaviors and reveal the unique ways in which mPFC DA encodes both the predictability and avoidability of aversive outcomes to enable rapid avoidance learning.

The DA system has a well-established role in associative learning, predominantly through studies examining striatal dopamine in reward-based tasks. Recent studies also implicate striatal DA in aversive learning^20,39–41^, in particular sustained avoidance behaviors supported by the tail of the striatum^42^. However, aversive processing robustly engages DA circuits projecting to the mPFC^16–18,43^, as aversive stimuli reliably drive increases in mPFC DA release^20,40,44,45^ and stimulation of prefrontal DA terminals biases behavior towards aversive behavioral responses^19^. Prior studies have demonstrated involvement of mPFC DA in the acquisition of contextual fear, as well as the expression—but notably not the acquisition—of cued fear^46–53^. Our results showing no requirement of mPFC DA for cue-shock associations are in line with these findings. Yet these previous studies primarily focused on fear conditioning or fully learned avoidance tasks. While the existing literature broadly implicates mPFC DA in aversive learning, our study demonstrates that rapidly learning to avoid aversive outcomes requires dynamic DA signaling in the mPFC. Learning may be driven primarily by a decrease in mPFC DA that signals when mice first take an action that causes omission of an aversive outcome.

Aspects of our studies were consistent with previous microdialysis studies that observed elevated mPFC DA as gerbils learned shuttle-box avoidance^24^, but we built on this work by revealing how event-linked mPFC DA activity changed throughout learning. We found that during both PMA and fear conditioning, mPFC DA levels were highest early in learning. While aversive foot shocks initially elicited large positive DA responses across all assays, in line with previous studies^18,40,54,55^, learning-related changes in the amplitude of these signals depended on the behavioral context. In particular, during fear memory retrieval shock omissions, there was a transient increase and rapid adaptation of mPFC DA, a unique pattern not observed in PMA or yoked conditioning. This supports previous work showing that VTA-mPFC projections regulate fear extinction learning^26,56–59^. In contrast to outcome-evoked DA signals, behaviorally-evoked DA transients were much smaller, and we observed subtle learning-related changes during platform entries. Together, these findings suggest that, rather than driving specific behaviors, bidirectional fluctuations in mPFC DA may promote learning-related circuit plasticity that is required to flexibly link predictive cues with adaptive actions required for avoidance.

Consistent with this interpretation, inhibiting VTA-mPFC DA slowed rather than prevented avoidance learning in PMA (Fig. 1H). However, our optogenetic manipulations are unlikely to have completely abolished VTA-mPFC DA activity. Thus, it is possible that a more complete suppression could fully block learning. Furthermore, VTA-mPFC DA terminal activity was necessary for learning avoidance strategies even in the absence of a motivational conflict but was not required for retrieval of previously learned avoidance behaviors. These results emphasize the role of VTA-mPFC DA in linking predictive cues to adaptive behaviors, rather than maintaining or enacting a learned behavioral strategy. While we did not observe a clear relationship between tone onset mPFC DA activity and PMA learning, there was a tone-induced response (Fig. 2G). Phasic stimulation of VTA-mPFC DA enhances the salience of a stimulus and improves subsequent discrimination learning^32^, so our optogenetic inhibition may have impaired learning by suppressing cue salience or blunting the contrast between shock-omission response and preceding activity during the tone.

At a cellular level, DA shapes prefrontal activity by influencing neuronal excitability and synaptic plasticity^60–62^, processes which may enable rapid avoidance learning. DA exerts these effects by acting primarily at D1 and D2 receptor classes, expressed by largely non-overlapping prefrontal cell types^63–66^. D1 -expressing mPFC projection neurons prominently project to cortical and thalamic targets^64,67^ and recent evidence shows that DA enhances signal-to-noise in mPFC projection neurons that encode aversive stimuli^19^. Such neurons play unique roles in threat avoidance learning^43,68^, so DA may shape activity in projection-specific mPFC populations to engage brain-wide networks required for avoidance learning. Future experiments should link activation of specific mPFC DA receptor subtypes with the patterns of circuit activity associated with avoidance learning. Altogether, our findings reveal that mPFC DA plays a distinct and essential role in avoidance learning by dynamically and bidirectionally encoding aversive outcomes and their omissions and using these representations to learn behavioral avoidance strategies. By dissociating the role of mPFC DA in active avoidance learning from its role in related behaviors, this study advances our understanding of the contribution of prefrontal DA to adaptive behavior. These findings also have significant implications for understanding disorders characterized by maladaptive avoidance, such as anxiety, post-traumatic stress disorder, and obsessive compulsive disorder^69,70^. Disturbances in prefrontal dopamine signaling could alter how predictive cues are linked to avoidance behaviors, contributing to pathological avoidance or excessive fear responses.

## Resource Availability

### Lead contact

Further information and requests for raw data and reagents will be fulfilled by the lead contacts, Dr. Laura DeNardo & Dr. Scott Wilke.

### Materials availability

This study did not generate new unique reagents.

### Data and code availability

- All data reported in this paper will be shared by the lead contact upon request.
- Any additional information required to reanalyze the data reported in this paper is available from the lead contact upon request.

## Acknowledgements

We thank Dr. Peter Balsam and Dr. Sotiris Masmanidis for helpful discussions of this work. This work was supported by a Klingenstein-Simons Fellowship in Neuroscience (LAD), a Whitehall Foundation Research Grant (LAD) and 1R01MH127214 (LAD), a Whitehall Foundation Research Grant (SAW), 1R01MH131858 (SAW), R01MH137461 (L.A.D, S.A.W), T32NS115753 (ZEZ), F32MH133387 (ZEZ), and T32NS115753 (TG).

## Author Contributions

LAD, SAW, ZEZ, MFG and TG conceptualized the experiments. ZEZ, MFG and MS performed experiments. ZEZ and TG analyzed data. LAD, SAW and ZEZ wrote the manuscript. All authors edited the manuscript.

## Declaration of Interests

The authors declare no competing interests.

## STAR METHODS

### Experimental Model and Subject Details

#### Animals

Adult male and female C57Bl6/J mice (JAX stock #000664) or tyrosine hydroxylase Cre (TH-Cre; line Fl12; www.gensat.org) mice were group housed (two to five per cage) and kept on a 12/12 h light/dark cycle (lights on 7AM to 7PM). Mice were sexed by examination of external genitalia at weaning. Animals received *ad libitum* food and water. All experiments were conducted in accordance with procedures established by the administrative panels on laboratory animal care at the University of California, Los Angeles.

### Method Details

#### Surgery

Mice were induced in 5% isoflurane in oxygen until loss of righting reflex and transferred to a stereotaxic apparatus where they were maintained under 2% isoflurane in oxygen. Mice were warmed with a circulating water heating pad throughout surgery and ophthalmic ointment was applied to the eyes. The head was shaved and prepped with three scrubs of alternating betadine and then 70% ethanol. Following a small skin incision, a dental drill was used to drill through the skull above the injection targets. A syringe pump (Kopf, 693A) with a Hamilton syringe was used for injections. Injections were delivered at a rate of 100 nl/min and the syringe was left in the brain for 10 min following injection. For optogenetics experiments, animals were injected with 500nL of AAV5-CAG-FLEX-JAWS-KGC-GFP-ER2 (Addgene #84445) or AAV5-CAG-FLEX-EGFP-WPRE (Addgene #51502) at a titer of 5×10^12^ vg/mL bilaterally into the VTA (AP: -2.9 & -2.6 mm, ML: +/- 0.45 mm, DV: -4.5 mm from bregma). Bilateral optic fibers (200 µm core, Newdoon) were implanted over the prelimbic (PL) area of the mPFC (AP: +1.8, ML: +/- 0.35, DV: -1.8 mm from bregma). For fiber photometry experiments, AAV5-hSyn-GRAB_DA2m (1×10^13^ vg/mL) was injected unilaterally into PL (AP: +1.8, ML: -0.35, DV: -2 mm from bregma) with an optic fiber (400 µm core, Doric) implanted 200 µm above it. For pain management mice received 5 mg/kg carprofen diluted in 0.9% saline subcutaneously. Mice received one injection during surgery and daily injections for 2 d following surgery. For optogenetic experiments, we waited at least four weeks after surgery before experimentation.

#### Behavior

##### Platform mediated avoidance

Three days prior to starting PMA, citric acid (2% w/v) was added to animals’ drinking water. This initiates a reduction in water consumption and increases performance on liquid reward tasks similar to forced water restriction^73^. Animals were then placed into a 20×22cm arena with metal rod flooring (Lafeyette Instruments) and a plexiglass platform covering one-quarter of the floor space. Opposite the platform was a reward port and reward pump (Campden Instruments). The reward port featured an infrared beam that triggered 25 µL of 10% sweetened condensed milk (Nestle) reward on a variable interval schedule (mean interval: 30 seconds). Once a day for two days, animals were placed in the arena and allowed to freely explore and consume reward for 45 minutes. The following day, PMA training began. For three days, animals were placed into the arena with a 180 second baseline period followed by delivery of a 30 second, 75 dB, 4kHz tone that co-terminated with a 2 second, 0.15 mA shock delivered to the floor (Lafeyette Instruments). Eight subsequent tone-shock pairings were delivered at randomly chosen intervals between 80 and 160 seconds. Training on PMA continued for two more days for a total of three days. Data presented in tone blocks are the average performance over three tones.

##### Fear conditioning and yoked control assay

For cued fear conditioning, animals were placed in the same arena as PMA without the platform or reward port. On training day, animals received three tones (4 kHz, 10s duration) to assess baseline freezing followed by twelve tones that were paired with a coterminal footshock identical to PMA. The following day (recall), animals were exposed to six tones (4 kHz, 30s duration) unpaired with shock. Data are presented as individual tones. For the yoked control experiment, the same arena as cued fear conditioning was used. Trial outcomes for individual subjects were matched to a PMA subject. On each of three days of training, subjects experienced nine tones (4 kHz, 30s duration) and based on the outcome of the matched PMA subject (avoid or shock), the shock was either omitted (for avoid outcome) or delivered (for shock outcome). Data presented in tone blocks are the average performance over three tones.

##### Optogenetics

Jaws activation was achieved via continuous red light (635 nm, 10 mW, Shanghai Laser and Optics Century) delivered through the entirety of the tone-shock periods. For PMA, laser illumination was concurrent with all the tones. For fear conditioning, this occurred on the first day (training).

##### Real-time place preference

After PMA, a subset of animals were placed in a real-time place preference (RTPP) assay. Animals were placed in a 68 x 23 cm chamber for 20 minutes. During the first ten minutes, the animals freely explored the arena to establish a baseline side preference. During the subsequent ten minutes, a closed loop video monitoring system (Bioviewer) tracked the location of the animal and triggered a 635 nm laser when the animal entered the “laser” side of the arena. The ratio of time spent in the “non-laser” side of the arena to the “laser” side of the arena was calculated for both baseline (laser off) and test (laser on) periods.

##### Fiber photometry

Recordings were performed using a commercial fiber photometry system (RZ10x, Tucker Davis Technologies) with two excitation wavelengths, 465 and 405 nm, modulated at 211 and 566 Hz respectively. Light was filtered and combined by a fluorescent mini cube (Doric Lenses). Emission was collected through the mini cube and focused onto a femtowatt photoreceiver (Newport, Model 2151). Samples were collected at 1017 Hz and demodulated by the RZ10x processor. Time stamps to synchronize experimental events and recordings were sent via TTLs to the RZ10x system via Arduino, controlled by custom MATLAB (MathWorks) code. For trial outcome specific analyses, trials were binned into groups of three as done earlier, using three subsequent trials of the same outcome to account for the stochastic nature of trial outcomes in PMA and yoked fear conditioning experiments.

#### Histology

Animals were deeply anesthetized with isoflurane and transcardially perfused with phosphate buffered saline (PBS), followed by 4% paraformaldehyde (PFA). Brains were removed and post-fixed overnight in 4% PFA before being moved to PBS. Brains were then sectioned via vibratome at 60 microns. To detect VTA axons expressing Jaws-GFP in the mPFC, anti-GFP immunohistochemistry was performed. Sections were incubated with chicken anti-GFP polyclonal antibody (1:2000, Aves Labs) overnight at 4℃. Sections were next rinsed and incubated with donkey anti-chicken AlexaFluor 488 conjugate (1:500; Jackson ImmunoResearch) at room temperature for 2 hr. Sections were washed before being mounted with DAPI. Images of fluorescent expression and implant targeting were taken using a Leica DM6 scanning microscope (Leica Microsystems) using a 10x objective.

#### Data Analysis

##### Behavioral Analysis

Overhead videos were collected (Teledyne FLIR, Chameleon 3) and animal positioning extracted via supervised deep learning networks^71^. Animal position information was transformed into freezing, platform entries, platform exits, and reward port entries using BehaviorDEPOT software^72^.

##### Fiber Photometry

Data were analyzed using a custom-written MATLAB pipeline. Raw recordings were downsampled 10x and the isosbestic signal was fit to the 465 nm signal using the polyfit MATLAB function. Signals were transformed into z-scores for tone, freezing, and platform periods using a baseline of -5 to -4 seconds relative to event onset and smoothed with a 25 frame moving average. An animal’s individual event responses were averaged to generate one trace per animal. Shock, avoid, and tone onset response z-scores were normalized across animals by using the mean response of the first tone period prior to shock delivery. Tone onset responses were calculated as the average value over the first 2 seconds of the tone. Shock/avoid responses were calculated as the average value over the 2 seconds following end of tone/shock. Platform and freezing responses were aligned to the onset or offset of the behavior. Average values of -2 to - 1.5 seconds, -0.5 to 0 seconds, and +1.5 to +2 seconds from the event were used as baseline, event, and post-event responses, respectively. Baseline values were set to 0 to calculate the changes in DA preceding and following the event.

In correlation plots, to account for different rates of learning, response metrics were correlated with trial outcomes following initial exposure to specific trial outcomes. Post-shock acquisition performance refers to fraction successful trials following the first three shock outcomes. Post-avoid acquisition performance refers to the fraction of successful trials following the first three avoid outcomes. For yoked and fear conditioned groups, avoid outcomes are replaced with shock-omission outcomes. Initial slope responses were calculated as the slope from the first block to the second block of that response type.

### Quantification and Statistical analysis

Data analysis were performed using custom scripts written in MATLAB. Statistical testing was performed in Graphpad Prism. All t-tests were performed as two-tailed. Error bars indicate SEM.

---

**Figure S1.**
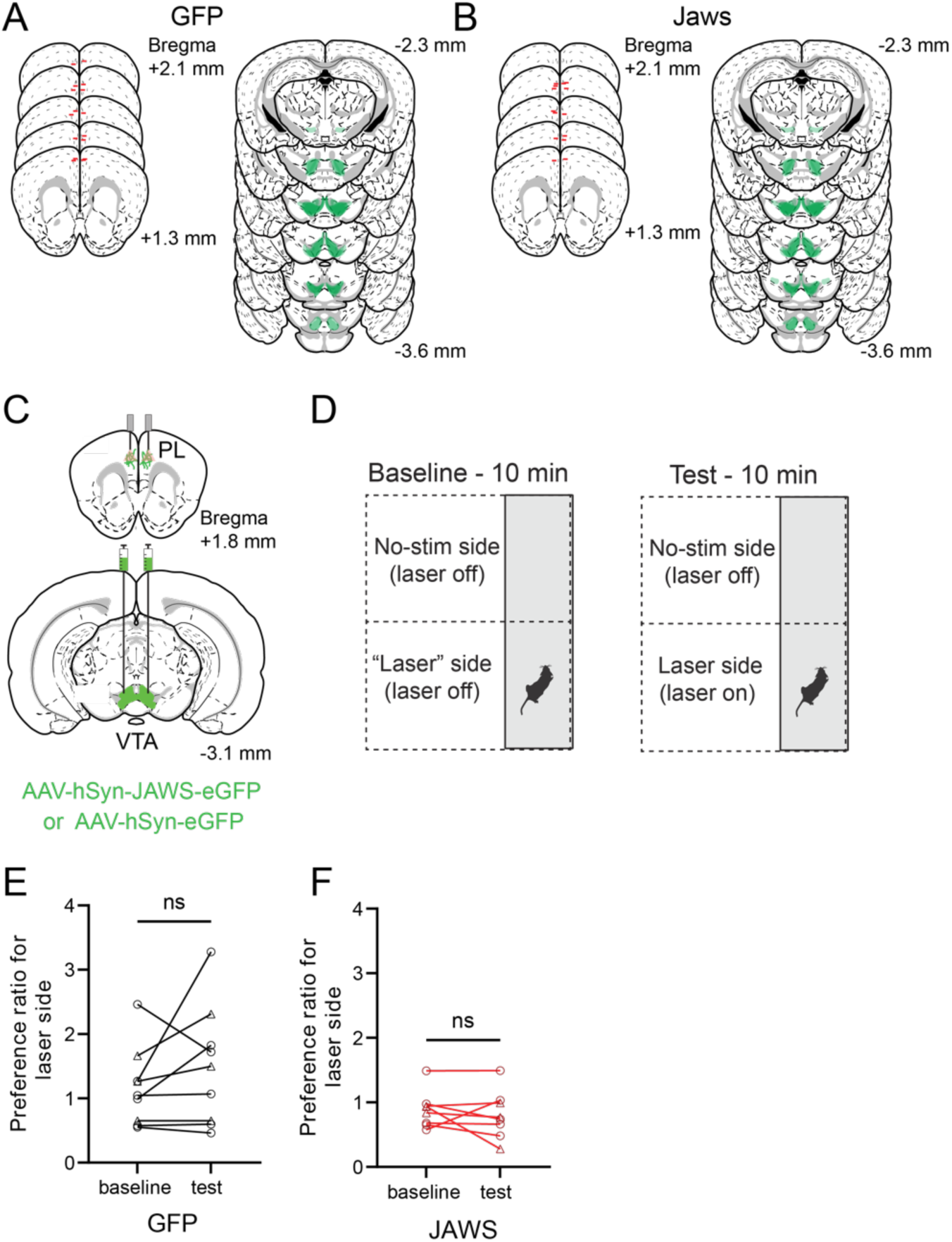
Tip placement and viral spread and real time place preference testing for optogenetic PMA experiments, Related to Figure 1. A. Tip placement (left, red lines) and viral spread (right, green) for GFP mice. B. Same as A for Jaws mice. C. Surgical strategy. D. Schematic of real-time place preference assay. E. Animal preference for “laser” side of the arena during baseline (laser off) period and during testing (laser on) period in GFP-expressing animals (paired t-test, P=0.2, n=9 mice). F. Same as E for Jaws-expressing animals (paired t-test, P=0.4, n=8 mice).

**Supplementary Figure 2.**
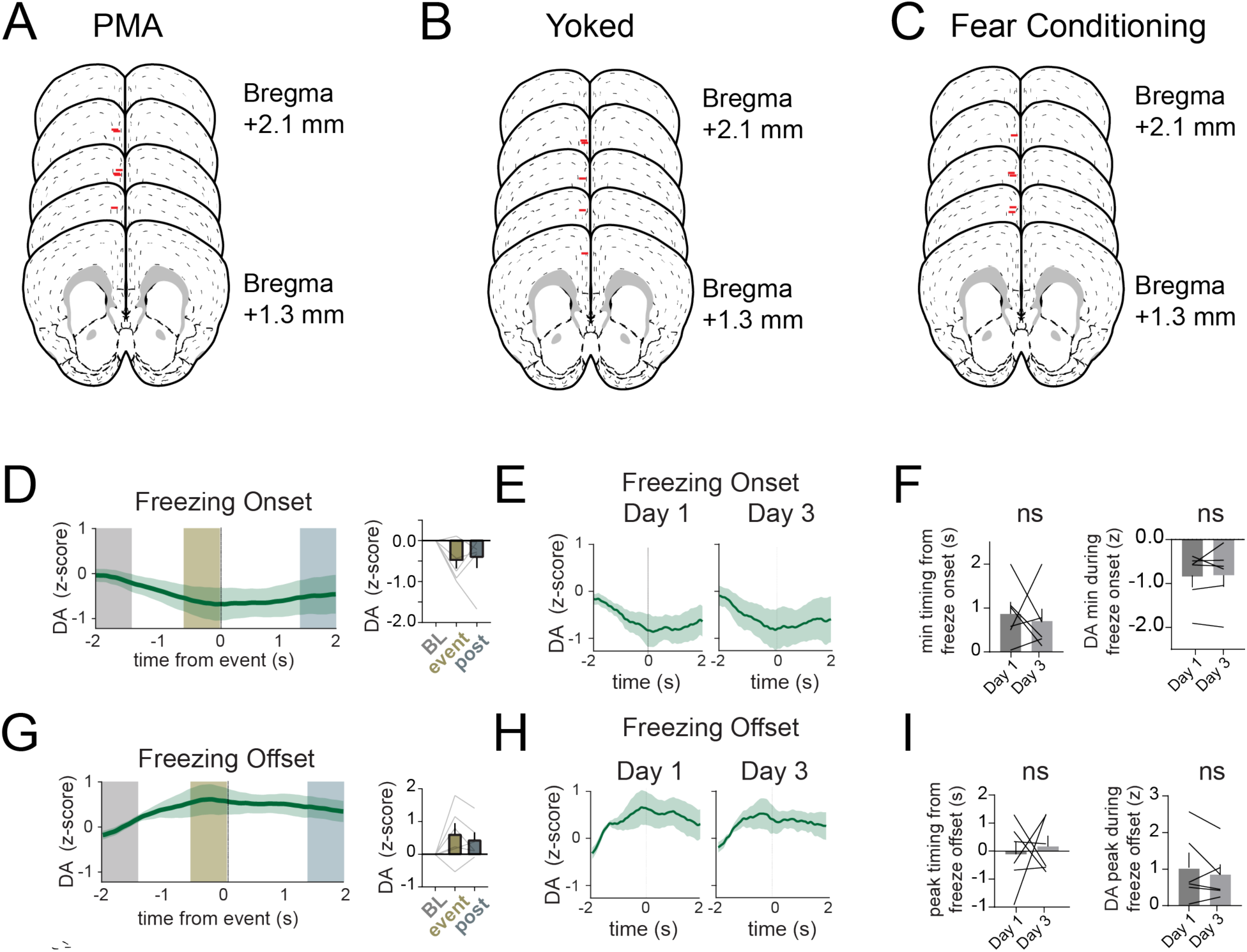
Tip placement for fiber photometry experiments and freezing-related DA dynamics across PMA learning, Related to Figure 2. A. Placement of tip (red lines) for fiber photometry recordings in PMA animals. B. Same as A for Yoked Fear Conditioning animals. C. Same as A for Fear Conditioning animals. D. DA dynamics during freezing onset (repeated measures one-way ANOVA F=2.8, P=0.1. n=6 mice). E. Comparison of freezing onset during Day 1 and Day 3 of PMA. F. (Left) Quantification of timing and (right) magnitude of freezing onset minimum values between Day 1 and Day 3. Timing (Wilcoxon signed rank test P=0.62, magnitude Mann Whitney P=0.93. n=6 mice). G. DA dynamics during freezing offset (repeated measures one-way ANOVA F=3.1, P=0.1). H. Comparison of freezing offset during Day 1 and Day 3 of PMA. I. (Left) Quantification of timing and (right) magnitude of freezing offset maximum values between Day 1 and Day 3. Timing (Wilcoxon signed rank test P=0.82, magnitude Mann Whitney P=0.86. n=6 mice). Graphs represent mean ± SEM.

**Supplementary Figure 3.**
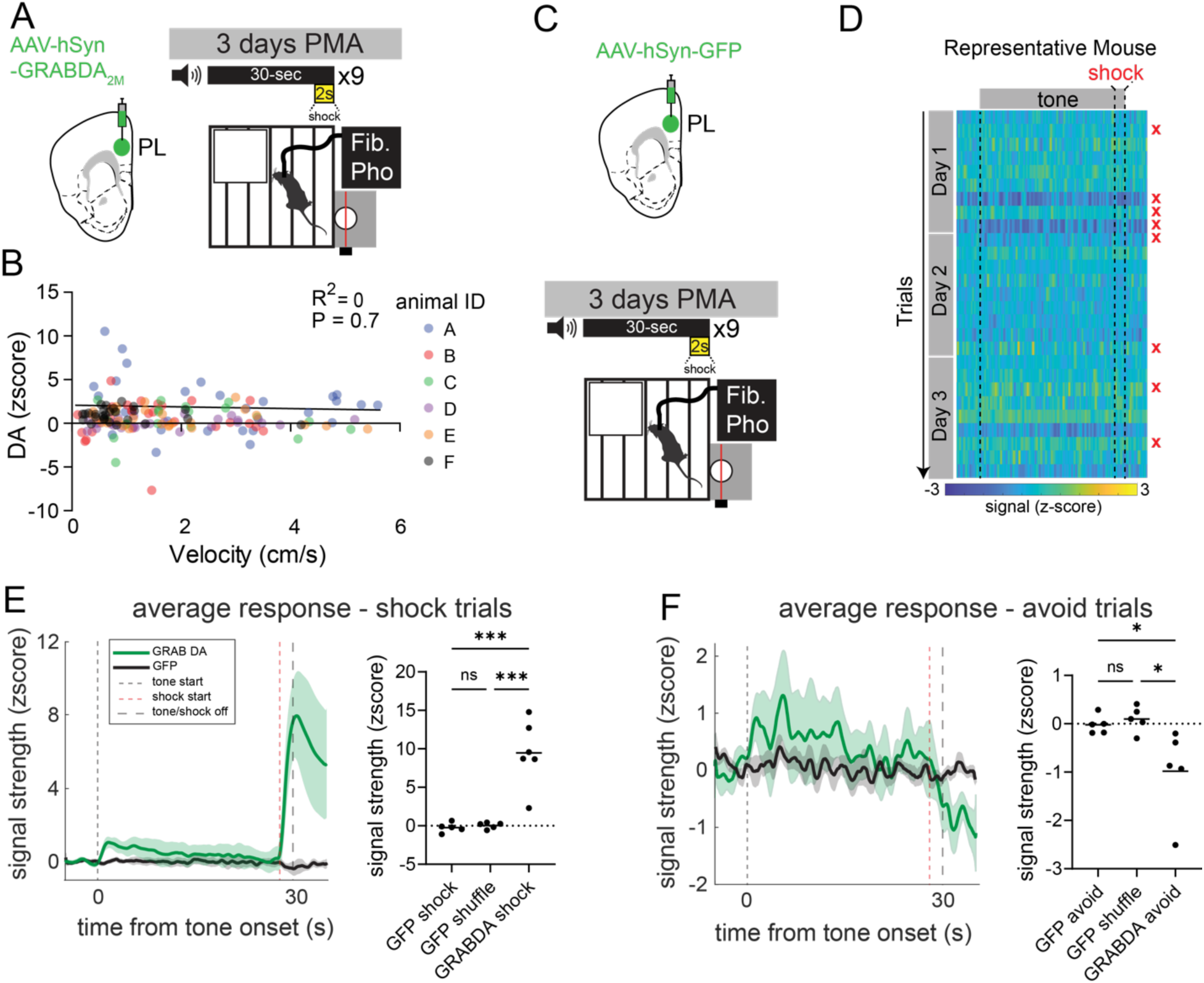
Dopamine responses are not caused by movement artifacts, Related to Figure 2. A. Surgical strategy to record GRABDA signal in PL (left); schematic of PMA (right). B. Velocity does not correlate with DA fluorescence. Data points are color coded by animal, with the average DA response and velocity at the start (first 3 seconds) of each tone (pearson’s R^2^ = 0, P=0.7). C. Surgical strategy to record fluorescent control (GFP) signal in PL (top); schematic of PMA (bottom). D. Representative heatmap of GFP signal recorded during PMA. E. (left) Averaged signal during shock trials for GFP animals (black) and GRAB DA animals (green). (right) Quantification of shock response in GFP animals, shuffled GFP signal, and GRAB DA animals (one-way ANOVA F(2,13)=22.6, P<0.0001. GFP shock vs GFP shuffle P=0.9, GFP shock vs GRABDA shock P=0.0002, GFP shuffle vs GRABDA shock P=0.0002. Tukey’s multiple comparisons tests. F. (left) Averaged signal during avoid trials for GFP animals (black) and GRAB DA animals (green). (right) Quantification of avoid response in GFP animals, shuffled GFP signal, and GRAB DA animals (one-way ANOVA F(2,12)=6.1, P=0.01. GFP shock vs GFP shuffle P=0.9, GFP shock vs GRABDA shock P=0.03, GFP shuffle vs GRABDA shock P=0.01). * P<0.05. *** P<0.001. Graphs represent mean ± SEM.

**Supplementary Figure 4.**
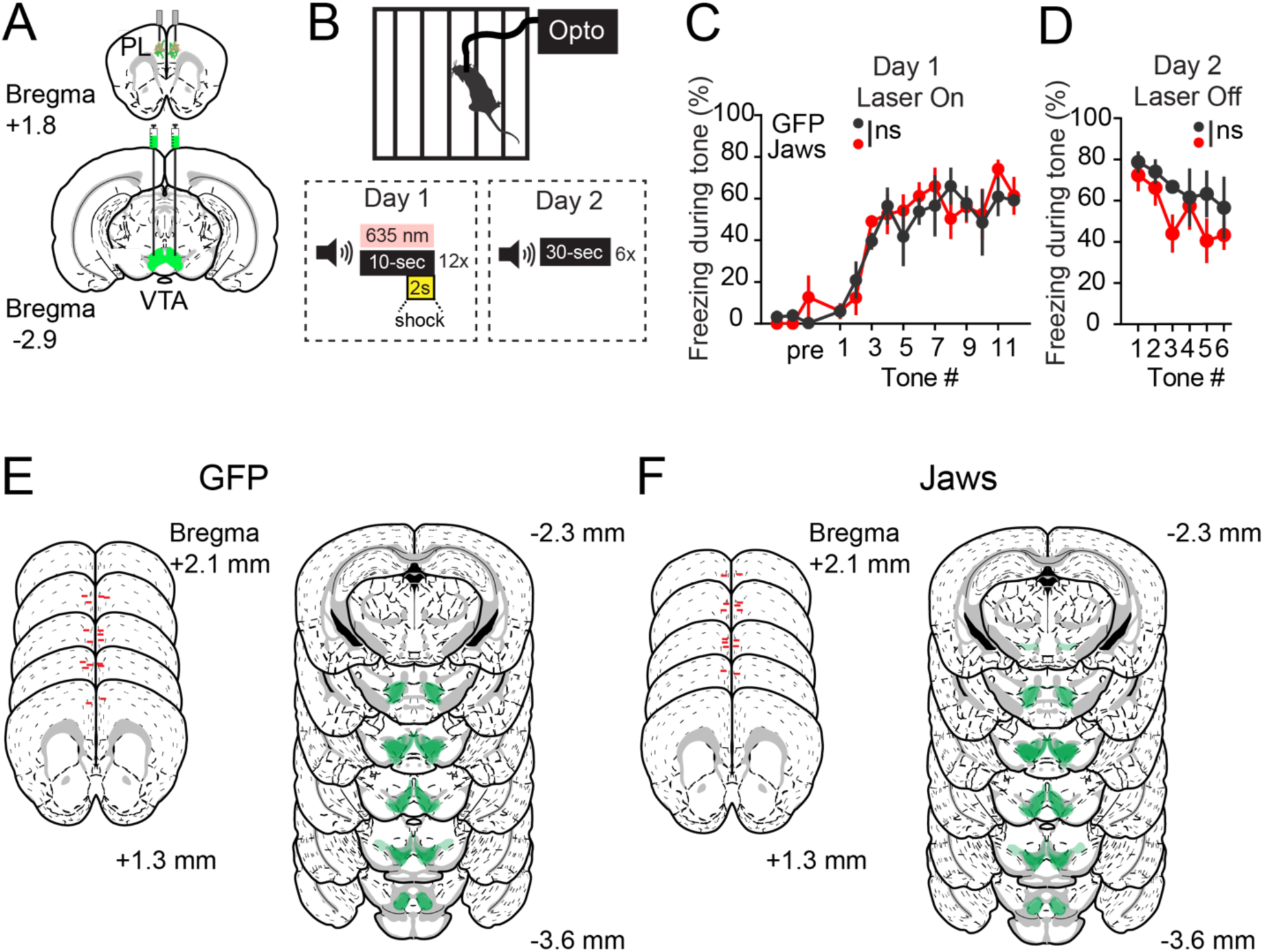
Optogenetic inhibition of mPFC VTA-DA terminals during fear conditioning, and viral and fiberoptic targeting for this experiment. Related to Figure 3. A. AAV injection and fiber placement for optogenetic inhibition of VTA-mPFC axon terminals. B. Experimental protocol. C. Freezing during fear conditioning (two-way repeated measures ANOVA (F_time_ (4.7, 33.3)=18.2, P<0.001; F_opsin_ (1, 7)=0.6, P=0.4; F_interaction_ (14, 98)=0.5, P=0.9, n=4 GFP, n=5 Jaws mice). D. Freezing during fear memory retrieval (two-way RM ANOVA (F_time_ (2.8, 20.2)=5.5, P=0.006; F_opsin_ (1, 7)=1.2, P=0.2; F_interaction_ (5, 35)=0.8, P=0.5, n=4 GFP, n=5 Jaws mice). E. Tip placement (top, red lines) and viral spread (bottom, green) for GFP mice. F. Same as A for Jaws mice. Graphs represent mean ± SEM.

## Notes

### Competing Interest Statement

The authors have declared no competing interest.

### Summary of Updates

Figures have been condensed to enhance conciseness of the article. New analyses are also included

## References

1. Mair, R. G., Francoeur, M. J., Krell, E. M. & Gibson, B. M. Where Actions Meet Outcomes: Medial Prefrontal Cortex, Central Thalamus, and the Basal Ganglia. Frontiers in Behavioral Neuroscience 16, (2022).

2. Matsumoto, K. & Tanaka, K. The role of the medial prefrontal cortex in achieving goals. Current Opinion in Neurobiology 14, 178–185 (2004).

3. DeNardo, L. A. et al. Temporal evolution of cortical ensembles promoting remote memory retrieval. Nature Neuroscience 22, 460–469 (2019).

4. Giustino, T. F. & Maren, S. The Role of the Medial Prefrontal Cortex in the Conditioning and Extinction of Fear. Frontiers in Behavioral Neuroscience 9, 1–20 (2015).

5. Zeidler, Z. & DeNardo, L. The Role of Prefrontal Ensembles in Memory Across Time: Time-Dependent Transformations of Prefrontal Memory Ensembles. in Engrams: A Window into the Memory Trace (eds. Gräff, J. & Ramirez, S.) 67–78 (Springer International Publishing, Cham, 2024). doi:10.1007/978-3-031-62983-9_5.

6. Euston, D. R., Gruber, A. J. & McNaughton, B. L. The Role of Medial Prefrontal Cortex in Memory and Decision Making. Neuron 76, 1057–1070 (2012).

7. Gabriel, C. J. et al. Transformations in prefrontal ensemble activity underlying rapid threat avoidance learning. Current Biology 35, 1128–1136.e4 (2025).

8. Diehl, M. M. et al. Active avoidance requires inhibitory signaling in the rodent prelimbic prefrontal cortex. eLife (2018) doi:10.7554/eLife.34657.

9. Diehl, M. M. et al. Divergent projections of the prelimbic cortex bidirectionally regulate active avoidance. eLife 9, 1–13 (2020).

10. Diehl, M. M., Bravo-Rivera, C. & Quirk, G. J. The study of active avoidance: A platform for discussion. Neuroscience and Biobehavioral Reviews 107, 229–237 (2019).

11. Moscarello, J. M. & LeDoux, J. E. Active avoidance learning requires prefrontal suppression of amygdala-mediated defensive reactions. Journal of Neuroscience (2013) doi:10.1523/JNEUROSCI.2596-12.2013.

12. Jercog, D. et al. Dynamical prefrontal population coding during defensive behaviours. Nature 595, 690–694 (2021).

13. Bravo-Rivera, C., Roman-Ortiz, C., Brignoni-Perez, E., Sotres-Bayon, F. & Quirk, G. J. Neural Structures Mediating Expression and Extinction of Platform-Mediated Avoidance. Journal of Neuroscience (2014) doi:10.1523/JNEUROSCI.0191-14.2014.

14. Capuzzo, G. & Floresco, S. B. Prelimbic and Infralimbic Prefrontal Regulation of Active and Inhibitory Avoidance and Reward-Seeking. Journal of Neuroscience 40, 4773–4787 (2020).

15. Weele, C. M. V., Siciliano, C. A. & Tye, K. M. Dopamine tunes prefrontal outputs to orchestrate aversive processing. Brain Research 1713, 16–31 (2019).

16. Lammel, S., Ion, D. I., Roeper, J. & Malenka, R. C. Projection-Specific Modulation of Dopamine Neuron Synapses by Aversive and Rewarding Stimuli. Neuron 70, 855–862 (2011).

17. Lammel, S. et al. Input-specific control of reward and aversion in the ventral tegmental area. Nature 491, 212–217 (2012).

18. Kim, C. K. et al. Simultaneous fast measurement of circuit dynamics at multiple sites across the mammalian brain. Nature methods 13, 325–328 (2016).

19. Vander Weele, C. M., et al. Dopamine enhances signal-to-noise ratio in cortical-brainstem encoding of aversive stimuli. Nature 563, 397–401 (2018).

20. Abercrombie, E. D., Keefe, K. A., DiFrischia, D. S. & Zigmond, M. J. Differential Effect of Stress on In Vivo Dopamine Release in Striatum, Nucleus Accumbens, and Medial Frontal Cortex. Journal of Neurochemistry 52, 1655–1658 (1989).

21. Del Arco, A., Park, J. & Moghaddam, B. Unanticipated Stressful and Rewarding Experiences Engage the Same Prefrontal Cortex and Ventral Tegmental Area Neuronal Populations. eNeuro 7, ENEURO.0029-20.2020 (2020).

22. Arnt, J. Pharmacological Specificity of Conditioned Avoidance Response Inhibition in Rats: Inhibition by Neuroleptics and Correlation to Dopamine Receptor Blockade. Acta Pharmacologica et Toxicologica 51, 321–329 (1982).

23. Wadenberg, M.-L., Ericson, E., Magnusson, O. & Ahlenius, S. Suppression of conditioned avoidance behavior by the local application of (−)sulpiride into the ventral, but not the dorsal, striatum of the rat. Biological Psychiatry 28, 297–307 (1990).

24. Stark, H., Bischof, A. & Scheich, H. Increase of extracellular dopamine in prefrontal cortex of gerbils during acquisition of the avoidance strategy in the shuttle-box. Neurosci Lett 264, 77–80 (1999).

25. Fibiger, H. C., Zis, A. P. & Phillips, A. G. Haloperidol-induced disruption of conditioned avoidance responding: attenuation by prior training or by anticholinergic drugs. Eur J Pharmacol 30, 309–314 (1975).

26. Luo, R. et al. A dopaminergic switch for fear to safety transitions. Nature Communications 9, 1–11 (2018).

27. Nader, K. & LeDoux, J. The dopaminergic modulation of fear: quinpirole impairs the recall of emotional memories in rats. Behav Neurosci 113, 152–165 (1999).

28. Greba, Q. & Kokkinidis, L. Peripheral and intraamygdalar administration of the dopamine D1 receptor antagonist SCH 23390 blocks fear-potentiated startle but not shock reactivity or the shock sensitization of acoustic startle. Behav Neurosci 114, 262–272 (2000).

29. Inoue, T., Izumi, T., Maki, Y., Muraki, I. & Koyama, T. Effect of the dopamine D(1/5) antagonist SCH 23390 on the acquisition of conditioned fear. Pharmacol Biochem Behav 66, 573–578 (2000).

30. Fadok, J. P., Dickerson, T. M. K. & Palmiter, R. D. Dopamine Is Necessary for Cue-Dependent Fear Conditioning. J Neurosci 29, 11089–11097 (2009).

31. Sun, F. et al. Next-generation GRAB sensors for monitoring dopaminergic activity in vivo. Nature Methods 17, 1156–1166 (2020).

32. Popescu, A. T., Zhou, M. R. & Poo, M.-M. Phasic dopamine release in the medial prefrontal cortex enhances stimulus discrimination. Proc Natl Acad Sci U S A 113, E3169–3176 (2016).

33. Hugues, S., Garcia, R. & Léna, I. Time course of extracellular catecholamine and glutamate levels in the rat medial prefrontal cortex during and after extinction of conditioned fear. Synapse 61, 933–937 (2007).

34. Yau, J. O.-Y. & McNally, G. P. The Activity of Ventral Tegmental Area Dopamine Neurons During Shock Omission Predicts Safety Learning. Behavioral Neuroscience 136, 276–284 (2022).

35. Jacobs, D. S. & Moghaddam, B. Prefrontal Cortex Representation of Learning of Punishment Probability During Reward-Motivated Actions. The Journal of Neuroscience 40, 5063–5077 (2020).

36. Park, J. & Moghaddam, B. Risk of punishment influences discrete and coordinated encoding of reward-guided actions by prefrontal cortex and VTA neurons. eLife 6, e30056 (2017).

37. Moorman, D. E. & Aston-Jones, G. Prefrontal neurons encode context-based response execution and inhibition in reward seeking and extinction. Proceedings of the National Academy of Sciences 112, 9472–9477 (2015).

38. McLaughlin, A. E., Diehl, G. W. & Redish, A. D. Potential roles of the rodent medial prefrontal cortex in conflict resolution between multiple decision-making systems. Int Rev Neurobiol 158, 249–281 (2021).

39. Thierry, A. M., Stinus, L., Blanc, G. & Glowinski, J. Some evidence for the existence of dopaminergic neurons in the rat cortex. Brain Res 50, 230–234 (1973).

40. Sorg, B. A. & Kalivas, P. W. Effects of cocaine and footshock stress on extracellular dopamine levels in the medial prefrontal cortex. Neuroscience 53, 695–703 (1993).

41. Mantz, J., Thierry, A. M. & Glowinski, J. Effect of noxious tail pinch on the discharge rate of mesocortical and mesolimbic dopamine neurons: selective activation of the mesocortical system. Brain Research 476, 377–381 (1989).

42. Menegas, W., Akiti, K., Amo, R., Uchida, N. & Watabe-Uchida, M. Dopamine neurons projecting to the posterior striatum reinforce avoidance of threatening stimuli. Nat Neurosci 21, 1421–1430 (2018).

43. Kajs, B. L., Loewke, A. C., Dorsch, J. M., Vinson, L. T. & Gunaydin, L. A. Divergent encoding of active avoidance behavior in corticostriatal and corticolimbic projections. Scientific Reports 12, 1–11 (2022).

44. Thierry, A. M., Tassin, J. P., Blanc, G. & Glowinski, J. Selective activation of the mesocortical DA system by stress. Nature 263, 242–244 (1976).

45. Sullivan, R. M. & Gratton, A. Relationships between stress-induced increases in medial prefrontal cortical dopamine and plasma corticosterone levels in rats: role of cerebral laterality. Neuroscience 83, 81–91 (1998).

46. Pezze, M. A., Bast, T. & Feldon, J. Significance of dopamine transmission in the rat medial prefrontal cortex for conditioned fear. Cereb Cortex 13, 371–380 (2003).

47. Vergara, M. D., Keller, V. N., Fuentealba, J. A. & Gysling, K. Activation of type 4 dopaminergic receptors in the prelimbic area of medial prefrontal cortex is necessary for the expression of innate fear behavior. Behavioural Brain Research 324, 130–137 (2017).

48. Díaz, F. C., Kramar, C. P., Hernandez, M. A. & Medina, J. H. Activation of D1/5 Dopamine Receptors in the Dorsal Medial Prefrontal Cortex Promotes Incubated-Like Aversive Responses. Frontiers in Behavioral Neuroscience 11, 209 (2017).

49. Feenstra, M. G. P., Teske, G., Botterblom, M. H. A. & De Bruin, J. P. C. Dopamine and noradrenaline release in the prefrontal cortex of rats during classical aversive and appetitive conditioning to a contextual stimulus: Interference by novelty effects. Neuroscience Letters (1999) doi:10.1016/S0304-3940(99)00601-1.

50. Wilkinson, L. S. et al. Dissociations in dopamine release in medial prefrontal cortex and ventral striatum during the acquisition and extinction of classical aversive conditioning in the rat. European Journal of Neuroscience 10, 1019–1026 (1998).

51. Feenstra, M. G. P. Dopamine and noradrenaline release in the prefrontal cortex in relation to unconditioned and conditioned stress and reward. in Progress in Brain Research vol. 126 133–163 (Elsevier, 2000).

52. Feenstra, M. G. P., Vogel, M., Botterblom, M. H. A., Joosten, R. N. J. M. A. & De Bruin, J. P. C. Dopamine and noradrenaline efflux in the rat prefrontal cortex after classical aversive conditioning to an auditory cue. European Journal of Neuroscience 13, 1051–1054 (2001).

53. Pezze, M. A., Marshall, H. J., Domonkos, A. & Cassaday, H. J. Effects of dopamine D1 modulation of the anterior cingulate cortex in a fear conditioning procedure. Progress in Neuro-Psychopharmacology and Biological Psychiatry 65, 60–67 (2016).

54. Abe, K. et al. Functional diversity of dopamine axons in prefrontal cortex during classical conditioning. eLife 12, RP91136 (2024).

55. Fadda, F. et al. Stress-induced increase in 3,4-dihydroxyphenylacetic acid (DOPAC) levels in the cerebral cortex and in n. accumbens: Reversal by diazepam. Life Sciences 23, 2219–2224 (1978).

56. Hitora-Imamura, N. et al. Prefrontal dopamine regulates fear reinstatement through the downregulation of extinction circuits. eLife 4, e08274 (2015).

57. Pfeiffer, U. J. & Fendt, M. Prefrontal dopamine D4 receptors are involved in encoding fear extinction. NeuroReport 17, (2006).

58. Zbukvic, I. C., Park, C. H. J., Ganella, D. E., Lawrence, A. J. & Kim, J. H. Prefrontal Dopaminergic Mechanisms of Extinction in Adolescence Compared to Adulthood in Rats. Frontiers in Behavioral Neuroscience 11, (2017).

59. Sartori, S. B. et al. Fear extinction rescuing effects of dopamine and L-DOPA in the ventromedial prefrontal cortex. Translational Psychiatry 14, 11 (2024).

60. Otani, S., Bai, J. & Blot, K. Dopaminergic modulation of synaptic plasticity in rat prefrontal neurons. Neurosci. Bull. 31, 183–190 (2015).

61. Buchta, W. C., Mahler, S. V., Harlan, B., Aston-Jones, G. S. & Riegel, A. C. Dopamine terminals from the ventral tegmental area gate intrinsic inhibition in the prefrontal cortex. Physiological Reports 5, e13198 (2017).

62. Chen, G., Greengard, P. & Yan, Z. Potentiation of NMDA receptor currents by dopamine D1 receptors in prefrontal cortex. Proceedings of the National Academy of Sciences 101, 2596–2600 (2004).

63. Floresco, S. B. & Magyar, O. Mesocortical dopamine modulation of executive functions: beyond working memory. Psychopharmacology 188, 567–585 (2006).

64. Anastasiades, P. G., Boada, C. & Carter, A. G. Cell-Type-Specific D1 Dopamine Receptor Modulation of Projection Neurons and Interneurons in the Prefrontal Cortex. Cereb Cortex 29, 3224–3242 (2019).

65. Anastasiades, P. G. & Carter, A. G. Circuit organization of the rodent medial prefrontal cortex. Trends in Neurosciences 44, 550–563 (2021).

66. Gee, S. et al. Synaptic Activity Unmasks Dopamine D2 Receptor Modulation of a Specific Class of Layer V Pyramidal Neurons in Prefrontal Cortex. J. Neurosci. 32, 4959 (2012).

67. Han, S.-W., Kim, Y.-C. & Narayanan, N. S. Projection targets of medial frontal D1DR-expressing neurons. Neuroscience Letters 655, 166–171 (2017).

68. Gongwer, M. W. et al. Brain-Wide Projections and Differential Encoding of Prefrontal Neuronal Classes Underlying Learned and Innate Threat Avoidance. J. Neurosci. 43, 5810–5830 (2023).

69. Kumar, M., Green, S. M. & Urs, N. M. Role of Cortical Dopamine circuits in regulating Striatal Dopamine dynamics during Reversal Learning. The FASEB Journal 36, (2022).

70. LeDoux, J. & Daw, N. D. Surviving threats: neural circuit and computational implications of a new taxonomy of defensive behaviour. Nature Reviews Neuroscience 19, 269–282 (2018).

71. Mathis, A. et al. DeepLabCut: markerless pose estimation of user-defined body parts with deep learning. Nature Neuroscience 21, (2018).

72. Gabriel, C. J. et al. Behavior DEPOT is a simple, flexible tool for automated behavioral detection based on marker less pose tracking. eLife 11, 1–33 (2022).

73. Anne E. Urai et al. Citric Acid Water as an Alternative to Water Restriction for High-Yield Mouse Behavior. eNeuro 8, ENEURO.0230-20.2020 (2021).

